# White Matter Lesion Asymmetry and Carotid Intraplaque Hemorrhage: A Novel Imaging Phenotype for Stroke Risk in Older Men

**DOI:** 10.64898/2026.01.27.702169

**Authors:** Nicholas Simard, Faraz Honarvar, Joshua Noronha, Adam Gibicar, Pascal Tyrrell, Alan Moody, M. Natasha Rajah, Sandra Black, Pejman Maralani, April Khademi

## Abstract

**Background:** While identifying high-risk carotid disease remains a significant clinical challenge, the specific role of carotid intraplaque hemorrhage (IPH) is poorly understood. Although IPH has been linked to white matter lesion (WML) burden, current assessments overlook the directional impact of plaque instability on the brain. This study sought to determine whether IPH is an independent driver of asymmetric WML pathology and evaluate if this phenotype can identify a high-risk demographic for TIA and stroke.

**Methods:** This multi-center retrospective study analyzed 264 participants (mean age 71.8 years) from the Canadian Atherosclerosis Imaging Network (2010–2015). Participants underwent 3T MRI to assess carotid IPH and WMLs. We quantified WMLs using a deep-learning pipeline to extract three biomarkers: volume (WML-ICV), intensity (WML-Intensity), and intensity ratio (WML-IR). The Asymmetry Index Measure (AIM) defined the inter-hemispheric log-ratio, while the association between IPH and AIM was examined using multivariable linear regression adjusted for age, sex, stenosis, and scanner manufacturer. A secondary composite outcome of TIA/stroke was analyzed via logistic regression to evaluate the interaction between IPH, age, and sex.

**Results:** While whole-brain WML burden did not significantly differ by IPH status (p > 0.60), IPH status was a robust independent predictor of hemispheric asymmetry (WML-ICV: p = 0.01; WML-Intensity, p = 0.01). Post-hoc analysis confirmed WML burden was significantly higher in IPH+ older males (≥ 70 years) compared to younger cohorts (p < 0.04). This older male subgroup also demonstrated 4.57-fold higher adjusted odds of TIA/stroke (p = 0.02) compared to other demographic subgroups (all p > 0.87).

**Conclusions:** Carotid IPH is independently associated with a rightward asymmetric WML phenotype not captured by global metrics. This imaging marker identifies a high-risk demographic of older males with a nearly five-fold increase in clinical events, suggesting that hemispheric-level analysis provides critical prognostic value for stroke risk stratification.

## 1 Introduction

Cerebrovascular disease (CVD) is a major life-threatening and high-morbidity disease [1]. Stroke, a type of CVD, affects more than 16.9 million people worldwide every year [1], making it the second-leading cause of death [2]. Given this profound negative impact on patients, families, and healthcare systems, CVD remains a critical area of investigation. Specifically, identifying and quantifying brain markers of CVD and its underlying causes is essential to enabling more appropriate and personalized treatment regimens to prevent irreversible neurological damage.

Magnetic resonance imaging (MRI) has been used to identify CVD features in the brain [3] such as white matter lesions (WMLs), stroke lesions, and lacunar infarcts [4, 5]. However, the pathogenesis of WMLs, especially in relation to carotid artery disease, remains unclear. Previous literature [6-11] has shown intraplaque hemorrhage (IPH), a common feature of atherosclerosis indicating plaque instability, may be associated with WML and other CVD features. For example, Altaf et al. [6] showed a significant interaction between carotid IPH and interhemispheric WML associated with a greater extent of ischemic brain damage [10]. Fan et al. [8] demonstrated IPH is associated with an elevated total burden of cerebral small vessel disease (SVD) in patients with carotid stenosis. Takaya et al. [9] showed carotid IPH is significantly associated with cerebrovascular ischemic events including stroke. Lastly, larger IPH volume has been associated with increased severity of stroke in patients with atherosclerosis [11].

Although WMLs are common markers of brain injury in CVD, prior studies have relied on subjective scoring and small single-center cohorts, which limit their generalizability and objectivity [3]. Few have explored whether hemispheric asymmetry in WMLs can reveal hidden patterns of brain vulnerability linked to IPH. Our study therefore proposes that WML asymmetry offers a more sensitive window into cerebrovascular health. Building on known sex and age differences in plaque genesis, we hypothesize that patients with IPH exhibit rightward WML asymmetry associated with adverse clinical outcomes. Using advanced segmentation tools and a novel Asymmetry Index Measure (AIM), we test this hypothesis in a diverse multi-center cohort, aiming to uncover a new imaging marker or phenotype of stroke risk.

## 2 Methods

### 2.1 Study Population

Data for this retrospective study was sourced from the Canadian Atherosclerosis Imaging Network (CAIN), a pan-Canadian study investigating CVD and carotid artery disease [12]. Participants were recruited across multiple Canadian academic medical centres between 2010 and 2015. The cohort include conventional demographic information (i.e. age, sex) and clinical follow-up data detailing outcomes including stroke and transient ischemic attack (TIA). Inclusion criteria for analyses were the availability of baseline Fluid-Attenuated Inversion Recovery (FLAIR) MRI and carotid Magnetic Resonance Intraplaque Hemorrhage (MRIPH) imaging. Exclusion criteria included presence of large cortical infarcts and imaging motion artifacts. Screening resulted in a final cohort of 264 subjects where the decision matrix can be seen in Figure 1A.

**Figure 1:**
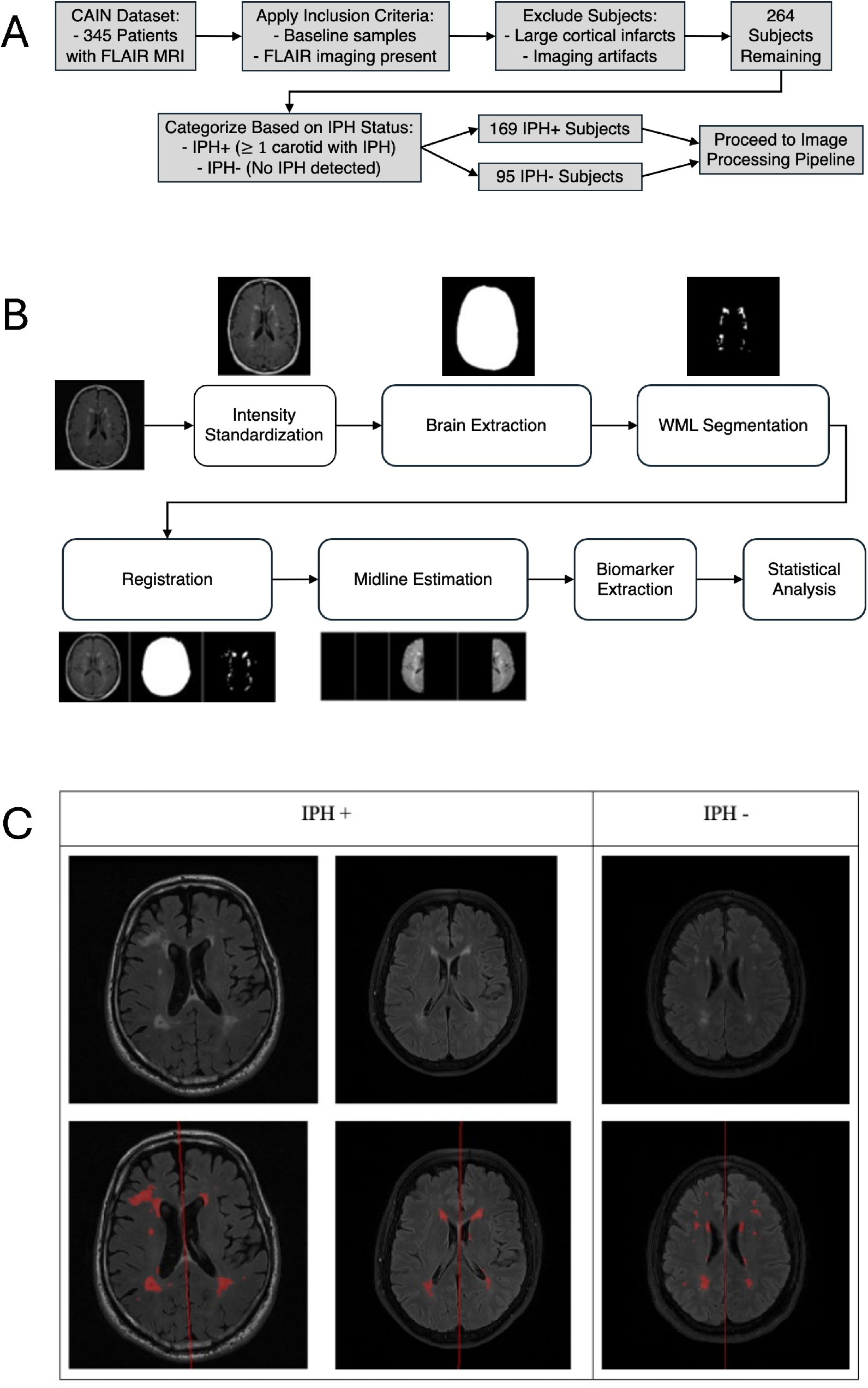
Patient selection flowchart and image processing pipeline. (A) Flowchart showing the participant selection process and inclusion criteria related to the CAIN dataset with the final cohort categorized into IPH+ and IPH– groups. (B) Image processing workflow detailing the atuomated pipeline used for WML quantification. (C) FLAIR MRI scans and automated segmentation outputs with the top row depicting original FLAIR images for two IPH+ subjects (left, middle) and one IPH-subject (right). The bottom row illustrates the processing results including segmented WMLs (red overlays) and the estimated hemispheric midline (red line).

### 2.2 Imaging Protocol

Structural brain imaging was acquired using 2D FLAIR MRI on 3T GE (Discovery MR750), Philips (Achieva), and Siemens (TrioTim, Skyra, and InteraMR) machines from 8 different centres. Acquisition parameters varied slightly across manufacturers GE (TR/TE/T1=9700/141/2200ms, 0.8954x0.8954x3mm^3^), Philips (TR/TE/T1=9000-11000/125/2800ms, 0.4286x0.4286x3mm^3^), and Siemens (TR/TE/T1=9000/117-119/2500ms, 1x1x3mm^3^). Carotid imaging included an MRIPH sequence (3D T1-weighted fat-suppressed fast field echo) for IPH identification and time-of-flight MR angiography (TOF-MRA) for stenosis measurements [13].

### 2.3 Image Analysis

High signal intensity areas in MRIPH identified atherosclerotic plaques [14] and quantified IPH for left and right carotid arteries. A plaque was classified as IPH-positive (IPH+) if signal intensity exceeded 150% of the adjacent muscle and IPH-negative (IPH-) otherwise [15], resulting in 169 IPH+ and 95 IPH-subjects. Comparatively, stenosis was measured via TOF-MRA and averaged across hemispheres for use as a technical covariate. FLAIR volumes were intensity standardized to mitigate multi-center effects [16]. Following brain extraction and intracranial volume (ICV) calculation, WMLs were segmented using a validated U-Net-variant deep learning model [17, 18]. Normal-appearing brain matter (NABM) was segmented by thresholding the standardized intensities to remove cerebrospinal fluid and hyperintense artifacts [16]. All volumes and masks were then registered to a FLAIR template space using ANTs SyN [19]. For asymmetry analysis, cerebral hemispheres were extracted from the registered data using a midline estimation method [20]. The entire image processing workflow is illustrated in Figure 1B with the WML segmentation shown in Figure 1C.

### 2.4 Quantification of WML Biomarkers

Three quantitative imaging biomarkers from the WML segmentation masks examined the relationship between disease asymmetry and IPH status. These biomarkers included the intracranial volume WML lesion load (WML-ICV), WML median intensity (WML-intensity), and the WML to NABM intensity ratio (WML-IR) [21]. Biomarkers were computed for the whole brain to assess total burden and in each cerebral hemisphere to facilitate asymmetry analysis. For each biomarker (*b)*, asymmetry was quantified using the novel Asymmetry Index Measure (AIM):

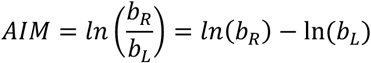

where b_*R*_ and b_*L*_ represent the right and left hemispheres. The AIM is positive (AIM > 0) if the features in the right hemisphere are higher (larger lesion load), and negative (AIM < 0) if the features in the left hemisphere are higher. The log-transformation was employed to handle skewed data distributions and to increase sensitivity to subtle inter-hemispheric differences in patients with mild-to-moderate WML disease [22].

### 2.5 Statistical Analysis

The primary analysis investigated whether IPH status independently predicts WML burden and asymmetry. Multivariable linear regression models were implemented for whole-brain biomarkers and corresponding AIM values, adjusting for known vascular risk factors and technical covariates. Sex was included to account for known differences in men being more likely to exhibit high-risk plaques, while Manufacturer controls for multi-vendor acquisition variability:

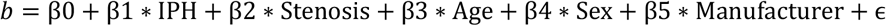

Hemispheric asymmetry was characterized by testing AIM values relative to a null hypothesis while ANCOVA was used to compare mean biomarkers across IPH laterality subgroups (negative, left, right, bilateral). Within the IPH+ cohort, a two-way ANOVA assessed the interactions between age (< 70 vs. ≥ 70) and sex (F/M), followed by Tukey’s honestly significant difference (HSD) post-hoc tests for pairwise comparisons.

The clinical relationships between WML pathology in the IPH+ cohort and adverse clinical events (TIA or stroke) were further examined using a logistic regression model. All statistical analyses were conducted leveraging packages in R (version 4.3.1) [24] and Python (version 3.13.5) [25]. Formal power calculations were not performed due to the retrospective design of this study.

## 3 Results

### 3.1 Patient Characteristics

The study included a total of 264 patients (mean age, 71.8 ± 7.7 years), where95 were IPH*−* and 169 were IPH+. Baseline demographics and clinical characteristics, including stenosis severity and scanner manufacturer distribution are summarized in Table S1. Clinical follow-up ranged from 0-30 months, with a median follow-up time of 23.9 months.

### 3.2 Whole Brain and Asymmetry Biomarkers

Multivariable linear regression revealed differences in WML burden between the IPH+ and IPH-cohorts with key statistical findings detailed in Table 1. Whole brain biomarkers (WML-ICV, WML-intensity, WML-IR) revealed no significant differences between IPH status cohorts (WML-ICV, p = 0.61; WML-Intensity, p = 0.63; WML-IR, p = 0.70), shown in Figure 2A. Moreover, AIM showed significant differences for WML-ICV, WML-Intensity, and WML-IR biomarkers between IPH+ and IPH-patients, shown in Figure 2B. In the IPH+ group, significant rightward asymmetry was observed in all three biomarkers (p = 0.01). The IPH-group had leftward asymmetry biomarkers that were not significantly different from 0 (WML-ICV, p = 0.11). No evidence was found for a difference in pulse pressure between IPH groups (p = 0.28).

**Table 1:**
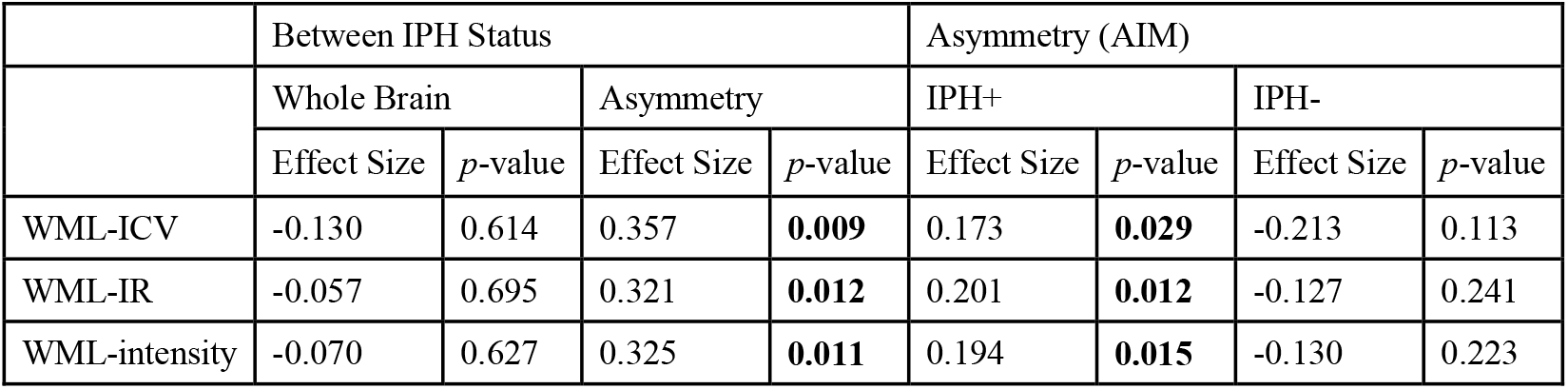
Statistical comparisons evaluating whole-brain burden and hemispheric asymmetry in IPH status cohorts. Effect sizes (Cohen’s d) and p-values are reported for inter-group differences in whole brain and AIM biomarkers, significantly deviating from zero or symmetric, within each IPH+ and IPH-cohorts. Bolded p-values indicate statistical significance (p < 0.05).

|  | Between IPH Status |  |  |  | Asymmetry (AIM) |  |  |  |
| --- | --- | --- | --- | --- | --- | --- | --- | --- |
|  | Whole Brain |  | Asymmetry |  | IPH+ |  | IPH- |  |
| | Effect Size | $p$-value | Effect Size | $p$-value | Effect Size | $p$-value | Effect Size | $p$-value |
| WML-ICV | -0.130 | 0.614 | 0.357 | **0.009** | 0.173 | **0.029** | -0.213 | 0.113 |
| WML-IR | -0.057 | 0.695 | 0.321 | **0.012** | 0.201 | **0.012** | -0.127 | 0.241 |
| WML-intensity | -0.070 | 0.627 | 0.325 | **0.011** | 0.194 | **0.015** | -0.130 | 0.223 |

**Figure 2:**
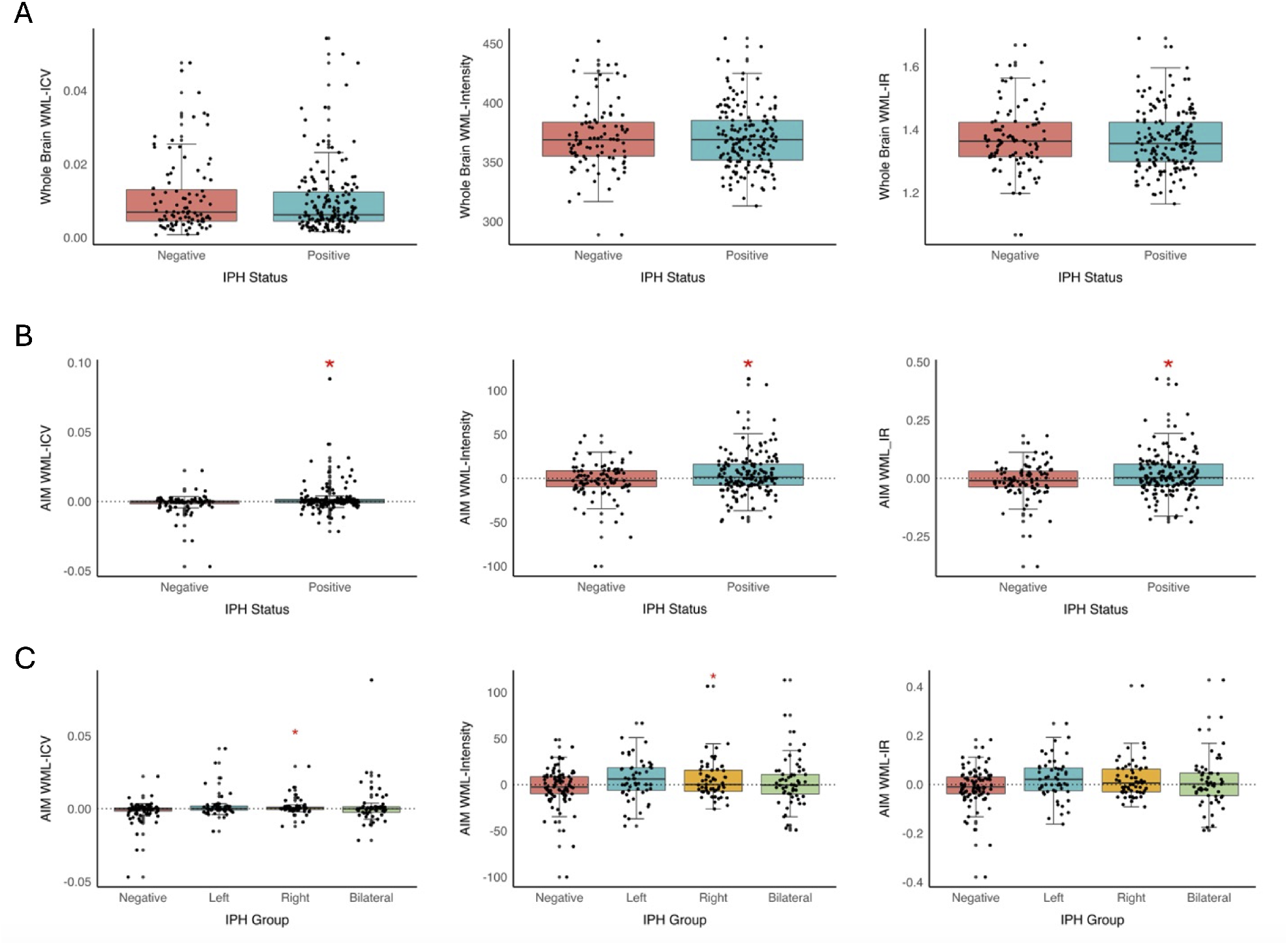
Comparison of whole-brain and AIM WML biomarkers by IPH status and laterality subgroups. (A) WML-ICV, WML-Intensity, and WML-IR biomarkers for whole brain in IPH+ and IPH-groups. (B) WML-ICV, WML-Intensity, and WML-IR biomarkers for AIM in IPH+ and IPH-groups. (C) Box plots of AIM biomarkers across IPH laterality subgroups. Dotted lines indicate the asymmetry threshold and a red asterisk indicates statistically significant rightward asymmetry in IPH+ patients.

### 3.3 Covariate Effects and ANCOVA

ANCOVA analysis, shown in Table 2, revealed that IPH status is a robust independent predictor for all metrics (p = 0.01). Conversely, IPH status showed no significant association with any whole-brain biomarkers (p > 0.57). Carotid stenosis was significantly associated with asymmetry in WML-Intensity (p = 0.01) and WML-IR (p = 0.01), but like IPH, did not predict whole-brain metrics (p > 0.23). Manufacturer effects were observed for asymmetry in WML-Intensity and WML-IR (p = 0.01). Notably, neither age nor sex were significant predictors of asymmetry (p > 0.19). To determine if these demographic factors acted as effect modifiers rather than independent predictors, a planned subgroup analysis was performed.

**Table 2:**
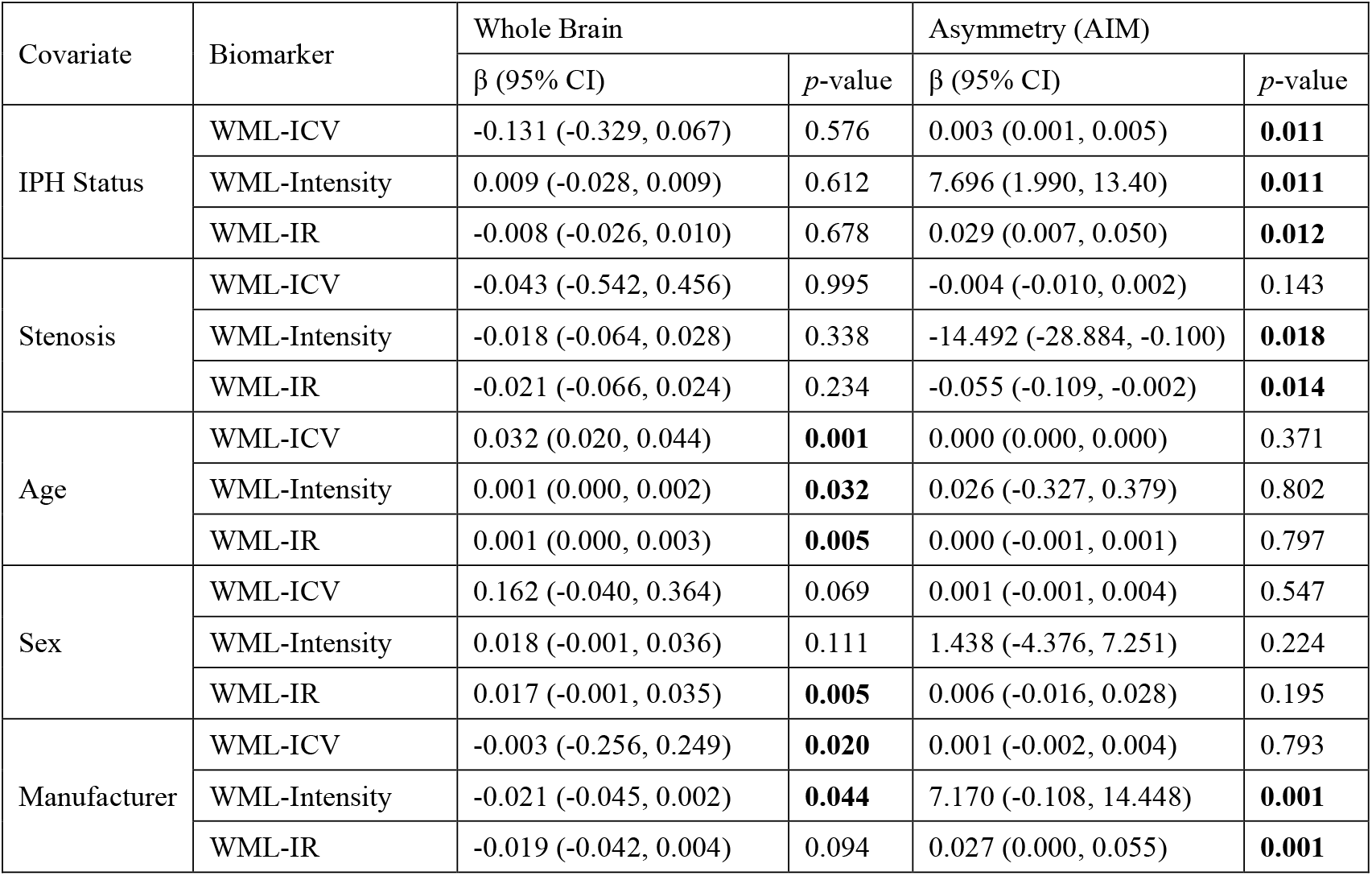
Linear regression analysis of whole-brain and AIM biomarkers associated with high-risk predictors. Coefficient estimates (β), 95% confidence intervals (CI), and p-values describe the independent influences of each predictor with WML pathology. Bolded p-values indicate statistical significance (p < 0.05), after adjusting for all other covariates in the model.

### 3.4 IPH+ Subgroup Analysis

AIM biomarkers were first examined across IPH subgroups (negative, left, right and bilateral), shown in Figure 2C. All IPH+ subgroups showed consistent rightward asymmetry, most pronounced in WML-ICV and WML-Intensity (p = 0.01). No evidence was found for asymmetry in the IPH-group (p = 0.11). Within the IPH+ cohort, a further subgroup analysis assessed the main effects of age (Younger <70 vs. Older ≥70) and sex (Female vs. Male) on whole brain and AIM biomarkers, shown in Table 3. A two-way ANOVA and post-hoc Tukey HSD test confirmed that WML burden was higher in the older male subgroup (≥ 70 years) across all AIM metrics compared to younger females (p < 0.02). When compared to younger males, older males maintained significantly higher asymmetry in WML-ICV and WML-IR (p < 0.02). Interestingly, older males showed significantly greater whole-brain WML-ICV burden than younger cohorts (p < 0.03). This demographic specific severity, highlighting older males as the most pathologically burdened subgroup, is shown in Figure 3.

**Table 3:** Post-hoc pairwise comparisons of whole-brain and AIM WML biomarkers grouped by age and sex using Tukey’s HSD test within the IPH+ cohort. Mean differences (Mean diff) represent the subtraction of the second group from the first, where positive values indicate higher burden in the first group. Adjusted p-values (P-adj) and 95% confidence intervals [lower, upper] are reported. Bolded P-adj values indicate statistical significance (p < 0.05), denoting distinct interactions within subgroups.

| Subgroups | Biomarker | Whole Brain |  |  | Asymmetry (AIM) |  |  |
| --- | --- | --- | --- | --- | --- | --- | --- |
|  |  | Mean diff | P-adj | 95% CI | Mean diff | P-adj | 95% CI |
| Older Female vs. Younger Female | WML-ICV | -0.003 | 0.455 | -0.009, 0.002 | -0.003 | 0.264 | -0.008, 0.001 |
|  | WML-Intensity | -7.598 | 0.698 | -25.702, 10.505 | -7.991 | 0.393 | -21.060, 5.078 |
|  | WML-IR | -0.025 | 0.753 | -0.090, 0.040 | -0.027 | 0.464 | -0.074, 0.020 |
| Older Female vs. Older Male | WML-ICV | 0.002 | 0.674 | -0.003, 0.006 | 0.002 | 0.482 | -0.002, 0.005 |
|  | WML-Intensity | 6.479 | 0.629 | -7.530, 20.487 | 4.734 | 0.622 | -5.378, 14.850 |
|  | WML-IR | 0.029 | 0.455 | -0.022, 0.079 | 0.022 | 0.394 | -0.014, 0.059 |
| Older Female vs. Younger Male | WML-ICV | -0.002 | 0.741 | -0.006, 0.003 | -0.002 | 0.631 | -0.005, 0.002 |
|  | WML-Intensity | -1.284 | 0.996 | -16.079, 13.511 | -3.090 | 0.878 | -13.771, 7.591 |
|  | WML-IR | -0.004 | 0.998 | -0.057, 0.049 | -0.011 | 0.894 | -0.049, 0.028 |
| Younger Female vs. Older Male | WML-ICV | 0.005 | **0.031** | 0.001, 0.010 | 0.005 | **0.003** | 0.001, 0.009 |
|  | WML-Intensity | 14.077 | 0.085 | -1.257, 29.410 | 12.725 | **0.017** | 1.656, 23.795 |
|  | WML-IR | 0.054 | 0.060 | -0.002, 0.109 | 0.049 | **0.009** | 0.009, 0.089 |
| Younger Female vs. Younger Male | WML-ICV | 0.001 | 0.886 | -0.004, 0.006 | 0.002 | 0.775 | -0.003, 0.006 |
|  | WML-Intensity | 6.314 | 0.739 | -9.741, 22.370 | 4.901 | 0.696 | -6.689, 16.492 |
|  | WML-IR | 0.021 | 0.775 | -0.036, 0.079 | 0.016 | 0.754 | -0.026, 0.058 |
| Older Male vs. Younger Male | WML-ICV | -0.004 | **0.034** | -0.007, -0.001 | -0.004 | **0.005** | -0.007, -0.001 |
|  | WML-Intensity | -7.763 | 0.282 | -19.000, 3.475 | -7.824 | 0.063 | -15.936, 0.288 |
|  | WML-IR | -0.032 | 0.165 | -0.073, 0.008 | -0.033 | **0.021** | -0.062, -0.004 |

**Figure 3:**
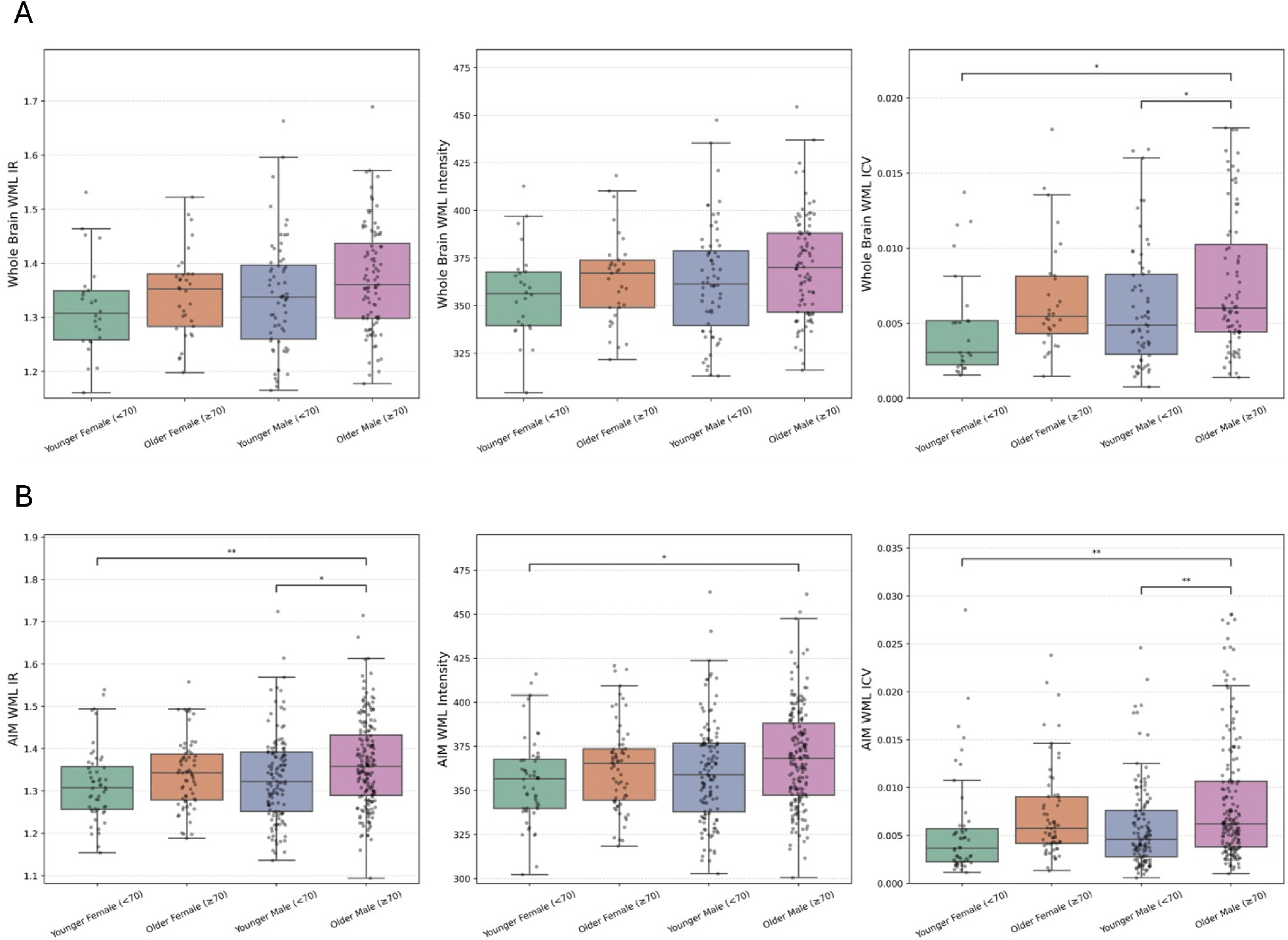
Subgroup analysis of whole-brain and AIMWML biomarkers in the IPH+ cohort. The top row illustrates whole-brain metrics while the bottom row corresponds to AIM metrics. Data is stratified by sex and age: Younger (< 70 years), Older (≥ 70 years), Female and Male subgroups. Horizontal brackets indicate statistically significant differences between subgroups (*p < 0.05, **p < 0.01).

### 3.5 IPH+ Group and Clinical Outcomes

For clinical outcome analysis, a logistic regression model evaluated the association between IPH status, age (Younger < 70 vs. Older ≥ 70) and sex (Female vs. Male) with clinical outcomes of TIA or stroke. To maximize statistical power given the limited stroke events (N=7) a composite outcome of TIA and Stroke was used and 2 subjects with an undefined IPH status were included. The model revealed that older males carried a significant 4.57-fold higher odds of TIA or stroke (OR = 4.57, p = 0.023). The event rate in this group was 12.5%, whereas no significant risk increase was observed in other demographic cohorts (p > 0.87). This suggests that IPH is a critical driver of clinical risk, particularly when stratified for age and sex.

## 4 Discussion

This multi-center study establishes a novel link between high-risk IPH features and asymmetric WMLs. This asymmetry, measured by WML-ICV, WML-Intensity, WML-IR, serve as sensitive biomarkers for systemic vascular instability extending beyond localized carotid disease.

We observed significant WML asymmetry in IPH+ subjects, consistent with literature suggesting the right hemisphere’s increased susceptibility to SVD due to blood supply distribution [26] and increased likelihood of injury and death [27, 28]. Notably, this rightward trend persisted across all IPH laterality groups, suggesting that IPH serves as a surrogate for poor overall vessel health rather than a purely local embolic source [8, 9, 11, 29]. Given the absence of whole-brain biomarker differences across groups and the connection to neurodegenerative states like Alzheimer’s disease [30], hemispheric asymmetry provides a highly sensitive and clinically valuable indicator of underlying vascular vulnerability.

ANCOVA results clarified the drivers of this lesion pathology. While both IPH and stenosis influenced lesion intensity (WML-Intensity/IR), WML-ICV was independently predicted by IPH status alone [7]. This suggests that while stenosis may contribute to tissue changes through chronic hypoperfusion, the volumetric expansion of lesions is more closely tied to the aggressive inflammatory nature of plaques. This is clinically significant because IPH can accelerate stenosis progression [31], increasing the likelihood of strokes [9], extracranial internal carotid artery stenosis [32], and reduced brain perfusion [33]. Furthermore, the insignificance of pulse pressure across IPH groups, confirms that our findings are independent of basic blood pressure differences and/or systemic hypertension alone.

A primary clinical contribution of this work is the identification of a high-risk phenotype. Subgroup analysis revealed that WML burden is not uniform, with older males (≥ 70 years) exhibiting significantly higher WML-ICV and WML-Intensity compared to younger cohorts. This imaging phenotype translated to a 4.57-fold higher odds ratio of TIA or stroke, transforming WML asymmetry into a critical risk stratification tool [34]. While current guidelines focus on stenotic degrees [35], our data suggests that rightward asymmetry on FLAIR MRI identifies patients who require earlier, more aggressive, pharmacological or lifestyle interventions [36]. The robustness of these findings is supported by our implemented pipeline and multi-center cohort [37]. Although scanner manufacturer influenced intensity, sensitivity analyses did not alter the rightward asymmetry findings, confirming a true physiological phenomenon rather than a technical bias.

Despite these strengths, several limitations remain. First, potential registration errors in the automated pipeline were statistically controlled but require close monitoring as they may influence observed manufacturer variability. Second, while a strong risk trend for older males persisted in the strictly IPH+ cohort, formal clinical significance was reached only when including subjects with undefined IPH status, requiring larger prospective studies to confirm subgroup effects. Finally, the study population and its demographics had limited generalizability for younger cohorts and detailed site-specific variables such as smoking and glycemic control were unavailable. Future research must validate these findings in independent datasets with more comprehensive vascular risk factor annotations.

Ultimately, this study demonstrates that hemispheric-level analysis is a critical addition to the clinical workup. WML asymmetry is not merely a precursor to global disease but represents a highly sensitive marker of underlying plaque instability and vascular vulnerability. While prior studies linked IPH with WML burden, none had previously examined directional asymmetry at this scale or with this level of automation. Our main finding shows that rightward WML asymmetry paired with the 4.57-fold higher odds of TIA or stroke in older males, identifies a specific high-risk phenotype that warrants immediate clinical attention. Therefore, WML asymmetry offers a novel imaging biomarker for early cerebrovascular vulnerability and should guide enhanced risk stratification and more aggressive clinical intervention in patients likely to harbor unstable carotid plaques.

## Non-standard Abbreviations and Acronyms

AIM: Asymmetry Index-Measure
CAIN: Canadian Atherosclerosis Imaging Network
CVD: Cerebrovascular Disease
CSF: Cerebrospinal Fluid
FLAIR: Fluid-Attenuated Inversion Recovery
ICV: Intracranial Volume
IPH: Intraplaque Hemorrhage
IR: Intensity Ratio
MRIPH: Magnetic Resonance Intraplaque Hemorrhage
NABM: Normal-Appearing Brain Matter
SVD: Small Vessel Disease
TIA: Transient Ischemic Attack
TOF-MRA: Time Of Flight Magnetic Resonance Angiography
WML: White Matter Lesions

## Acknowledgements

The authors would like to thank the members of the Image Analysis in Medicine Lab (IAMLAB) at the Toronto Metropolitan University for supporting the project.

## Authorship

A.M., P.M., and A.G. conceived and planned the project. N.S., F.H., A.G., and J.N. carried out the sample preparation, data gathering, and analysis. A.K., A.M., P.T., P.M., S.B., N.S., and F.H. contributed to the interpretation of the results. N.S., F.H., and J.N. wrote the final manuscript. All authors provided critical feedback and helped shape the research, analysis, and manuscript.

## Supplemental Material

Table S1

STROBE Checklist

